# Synthetic Horizontal Gene Transfer for Ecosystem Restoration

**DOI:** 10.1101/2025.09.23.678013

**Authors:** Victor Maull, Guim Aguadé-Gorgorió, Victor de Lorenzo, Ricard Solé

**Affiliations:** ICREA-Complex Systems Lab, University Pompeu Fabra, Dr Aiguader 80, 08003 Barcelona; Institut de Biologia Evolutiva, CSIC-UPF, Pg Maritim de la Barceloneta 37, 08003 Barcelona; ISEM, University Montpellier, CNRS, IRD, Montpellier, France; Centro Nacional de Biotecnologia, CSIC, Madrid, Spain; Santa Fe Institute, 1399 Hyde Park Road, Santa Fe NM 87501, USA

**Keywords:** Synthetic Biology, Ecological Engineering, climate change, catastrophic shifts, mutualism

## Abstract

Restoring endangered ecosystems has become a pressing issue as the effects of global warming continue to harm existing communities. Over the last decade, several studies have suggested the use of synthetic biology as a tool to protect these biodiverse communities by augmenting their functionality. A critical example concerns soil microbiome communities in drylands, where increasing water retention by some of the constituent species could effectively protect the ecosystem from abrupt degradation. However, how to effectively deploy a functional synthetic construct that can scale its impact to the community level remains an open question. Recent experimental research has designed recombinant gene plasmids with the capacity to horizontally transfer across soil microbial communities. Here, we explore the impacts of synthetic horizontal gene transfer in models of ecological consortia. By coupling consumer-resource dynamics with multispecies gene transfer, our work identifies the design conditions that promote biodiversity while regulating gene propagation and the spread of engineered organisms.

## I. INTRODUCTION

During the past two centuries since the Industrial Revolution, anthropogenic pressures, including accelerating climate change, pollution, intensive agriculture, grazing, and soil erosion, compounded by widespread habitat loss and biodiversity, have placed the stability of the planet in jeopardy. Many of these trends continue to accelerate, and models project multiple scenarios of ecosystem collapse in the coming decades (Berdugo et al., 2020; Lenton et al., 2008; Levin, 2000; Scheffer, 2020; Solé and Levin, 2022). In response, global efforts have increasingly focused on sustainable land and resource management, biodiversity conservation, renewable energy transitions, and sustainable development (Rockström et al., 2024). However, these measures alone may be insufficient, as multiple processes in the Earth’s system are rapidly approaching or exceeding the planetary boundaries that define a safe operating space for humanity (Steffen et al., 2015).

Beyond mitigation and conservation, engineered intervention strategies have been proposed, such as geoengineering approaches for carbon removal or large-scale afforestation. These top-down methods offer more controllable results, but often come with high costs, scalability constraints, and uncertain ecological side effects (Society, 2009). In contrast, an alternative approach draws on the intrinsic capability of living systems themselves. Based on synthetic biology, this bottom-up restoration strategy proposes the design or modification of microbial communities to restore ecosystem functionality or confer resilience under environmental stress (Cases and de Lorenzo, 2005; Conde-Pueyo et al., 2020; de Lorenzo, 2008, 2017; de Lorenzo et al., 2016; Solé, 2015; Vidiella et al., 2020). Examples include the restoration of coral reefs (van Elsas et al., 2012) and soils (Jansson et al., 2023), the improvement of the dryland microbiota (Maestre et al., 2017), and broader frameworks for ecological bioengineering (de Lorenzo, 2022). Although diverse in mechanism and context, these strategies coalesce around a shared goal: employing biological agents to steer ecosystems back toward desirable, stable states or toward new, resilient configurations, see Fig. 1a. In this paper, we build on these concepts by advancing a synthetic biology–based intervention scenario, with engineered microbes serving as restoration agents capable of sustainably improving ecosystem recovery.

**FIG. 1.**
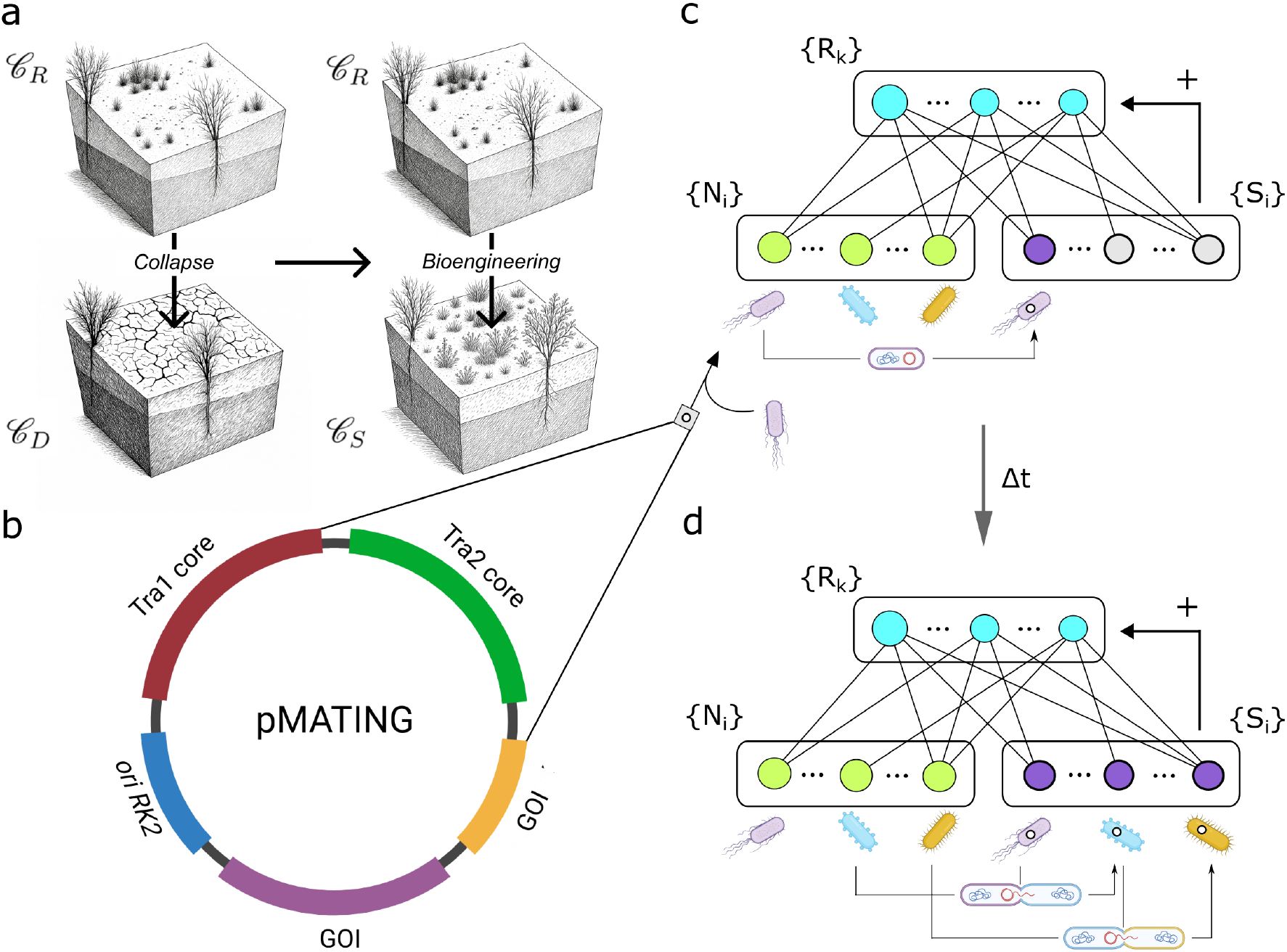
Engineering ecological networks using Horizontal Gene Transfer. (a) Ecosystem trajectories. An endangered ecosystem (𝒞_*R*_) may either collapse (𝒞_*D*_) or be shifted toward recovery (𝒞_*S*_) through bioengineering. (b) The “pMATING” system enables propagation of recombinant genes via horizontal transfer in microbiomes. A gene of interest (GOI) is inserted into a bacterial host, which then spreads the plasmid via conjugation. (c) The model approach is based on a resource–consumer network model. Resources (*R*_*k*_) are consumed by native (*N*_*i*_) and synthetic (*S*_*i*_) plasmid carrier species. Initially, only the re-introduced host in (*S*_*i*_) carries the plasmid. (d) After Δ*t*, horizontal transfer occurs, leading to competition and GOI expression, which enhances resource stability. This mechanism could promote ecosystem recovery.

The starting point is the identification of naturally occurring or laboratory-engineered biological materials – such as microbial strains or consortia– that possess superior catalytic abilities. These may range from the degradation of recalcitrant or toxic chemicals to fixation with *CO*_2_ and *N*_2_, to capture residual humidity or tolerating soil alkalinization after fires. In addition, modern genetic techniques make it possible to design and program microorganisms *a’ la carte*, combining traits within a single agent that are otherwise scattered in nature. This opens the door to an unprecedented repertoire of biological activities to mitigate the harmful impacts of urban and industrial processes. However, the challenge lies not only in discovering or enhancing (whether recombinantly or not) such active biological agents but also in developing effective strategies for their large-scale delivery and persistence. By analogy to the administration of drugs to the human body, this problem has been appropriately termed Environmental Galenics (de Lorenzo, 2022). In other words, once we have an activity of environmental interest in hand, how can we deliver it extensively to target ecosystems safely and effectively?

A key challenge for all bioengineering strategies is ensuring that laboratory-made constructs spread effectively, ideally scaling with community size. In this respect, it is highly informative to leverage how microbial communities naturally acquire new biodegradation or resistance traits to cope with environmental stress. Typically, this occurs when one member of the community evolves a solution to a specific challenge, and that DNA-encoded innovation is subsequently propagated, sometimes very rapidly, to neighboring organisms and other community members. Therefore, horizontal gene transfer (HGT) involves all transfers of genetic material from one cell to another. HGT has traditionally been described through three distinct mechanisms: conjugation, transformation, and transduction. Conjugation, in particular, requires direct contact between donor and recipient cells via a pilus, through which genetic material is transferred, and appears to be the most promising short-term strategy for DNA transfer to a wide range of potential bacterial recipients (Arnold et al., 2021; de Lorenzo, 2022; Soucy et al., 2015). This principle is particularly evident in the spread of antibiotic resistance. Although this phenomenon raises major concerns in the biomedical field, it also reveals the existence of highly efficient DNA transmission channels that enable the rapid dissemination of beneficial traits across communities.

Unfortunately, the strong association between HGT and antibiotic resistance has often obscured the enormous opportunity to repurpose these very same mechanisms for positive outcomes. By harnessing conjugation and related processes through synthetic biology, it becomes possible to deliberately propagate advantageous activities; a kind of targeted enhancement of the catalytic capacities of microbes within a given ecosystem. This is particularly relevant in the context of the central role that microbial diversity plays in a wide range of ecological scenarios (Banerjee and Van Der Heijden, 2023). This includes diversity as a barrier against invaders and pathogens (Case, 1990; van Elsas et al., 2012), as a driver of multifunctionality (Delgado-Baquerizo et al., 2016a), climate mitigation (Beattie et al., 2025), and as an in-trinsic component of engineering strategies where it acts as a firewall (Maull and Solé, 2024).

In this context, the work of Aparicio et al. (2022) is especially noteworthy (Aparicio et al., 2022). This work has experimentally demonstrated the feasibility of engineering hyperpromiscuous plasmids (Fig. 1b) in laboratory donor strains, enabling the in situ transfer of combinatorial gene sets to diverse soil bacteria. Although the study was limited to a proof-of-concept model system, the results strongly suggest that scaling up this strategy with vectors carrying functions of environmental relevance could provide powerful tools for restoring ecosystems damaged by chemical pollutants and atmospheric stressors. Could these engineering approaches be used to restore ecosystems?

In order to address the previous questions, mathematical modeling efforts can help guide experimental design and deepen mechanistic understanding of ecological tipping points and the engineering to restore (Solé et al., 2015, 2018; Vidiella and Solé, 2022). In addition, modeling diverse communities can help predict the potential outcomes of HGT-based intervention scenarios. This paper aims to support such approaches by proposing a minimal model that incorporates three key ecological features, namely:

1. Multi-species interactions, to reflect community-level dynamics and account for the role of biodiversity in resisting ecosystem collapse and providing ecosystem services and reliability through functional redundancy. (Case, 1990; Delgado-Baquerizo et al., 2016b; Hooper et al., 2005; Naeem and Wright, 2003).
2. Environment-community feedbacks, with niche construction (Kylafis and Loreau, 2010) as a central concept, modeled using extensions of the Lotka-Volterra equations (Chesson, 1990; MacArthur, 1970).
3. Gene spread as an epidemiological process, based on classical models of HGT (Kermack and McKendrick, 1927; Rose, 1983; Tremblay and Rose, 1985; Zhu et al., 2024).

This approach enables the integration of desired functions at the multispecies level, overcoming the limitations of single-species engineering. This paper aims to examine the potential effects and robustness of this strategy. As our model shows, HGT spreading of a functionally beneficial construct can enhance both the efficiency of the intervention and its community-level impacts while having no cascade effects nor other unintended consequences.

## II. METHODS

Here, a two-layered network *ξ* of interaction between the extant species of a given community and the available resources, see Fig. 1c. In parallel, a potential, yet non-existent, synthetic community shares the same *ξ* interaction context. The first synthetic resident would be introduced through an initial anthropogenic action: the addition of a synthetic species, which essentially comprises a modified native species that harbors a highly transmissible plasmid, i. e. plasmid pMATING, described by (Aparicio et al., 2022). See Fig. 1b. Once the first synthetic is introduced, the propagation of the recombinant genes will spread across the extant community by HGT, see Fig. 1d. Interestingly, the synthetic genetic material accommodates a gene of interest section (GOI) for delivering heterologous genes to wild-type mating partners. Such GOI can potentially induce specific functionalities that would benefit the endangered community.

In a previous work (Maull and Solé, 2024), we studied how slowing down the depletion of shared resources leads to a rise in biomass and biodiversity. Additionally, we highlight the seamless integration of synthetic species into the ecological network, accompanied by minor yet positive shifts in the population size of the existing community. One particular scenario is outlined in (Maull and Solé, 2024). A synthetic species capable of producing an exopolymer could help mitigate water loss, thereby positively influencing the other community members. For example, the production of a simple exopolymer, such as hyaluronic acid (Sze et al., 2016; Widner et al., 2005), might involve a cassette of genes encoded within the gene of interest (GOI) section of a pMATING plasmid.

We tackle mathematically this problem by building a set of three differential equations incorporating (1) inorganic resource availability, denoted by *R*_*k*_, where *k* = 1, 2, …*m*, (2) a set of resident species, denoted by *N*_*i*_, where *i* = 1, 2, …*n*, and (3) the synthetic pool, denoted by *S*_*i*_, with the same maximum size. For clarity, a diagram of the model is shown in Fig. 2.

**FIG. 2.**
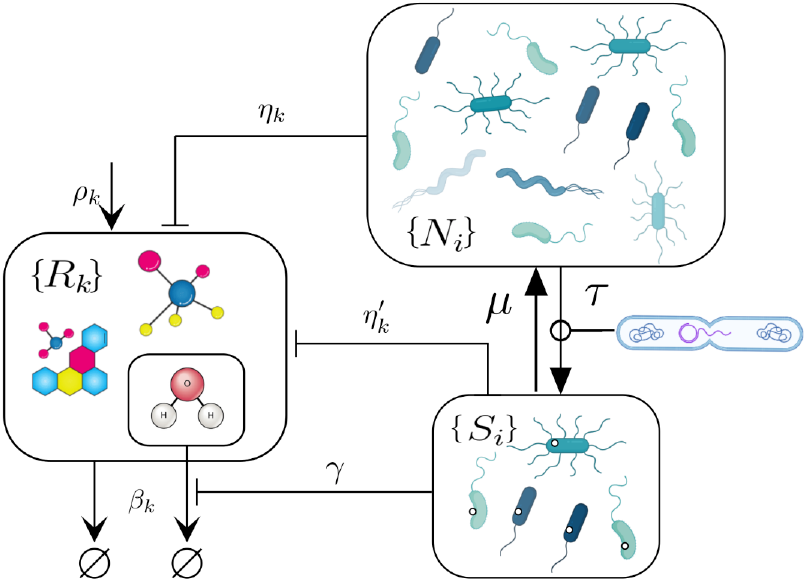
The multi-resource, multi-species framework. The extant community resides the upper box as {*N*_*i*_ }, growing while consuming resources from the left compartment {*R*_*k*_} at a given rate. Upon the introduction of the first synthetic inoculum, horizontal gene transfer (HGT) enables plasmid transmission across the community, causing a fraction of species to transition into the synthetic compartment {*S*_*i*_ } at a rate *τ* . These synthetic species also consume resources from the same pool. Plasmid loss occurs at a rate *µ*, leaking back to {*N*_*i*_ }. Additionally, the synthetic community {*S*_*i*_} affects {*R*_*k*_} decay rate *β*_*k*_.

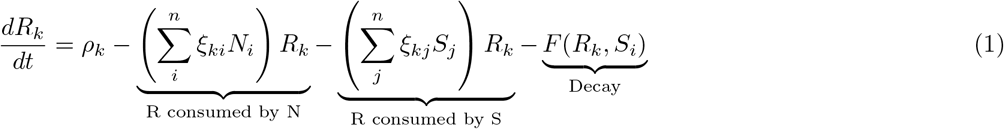

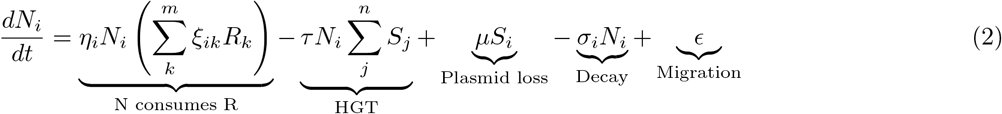

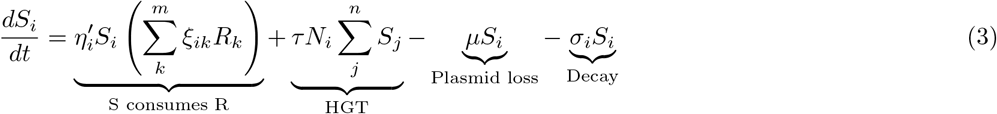

The resources equation has four main parts. The intrinsic, resource-dependent growth rate is represented by *ρ*_*k*_ ∈ [0, 1], the consumption term by *N*_*i*_, the consumption term by *S*_*i*_, and a decay term *F* (*R*_*k*_, *S*_*i*_) that entangles the synthetic functionality within which the mechanism is detailed separately below.

The equation of extant species has five main parts. The growth rate *η*_*i*_ ∈ [0, 1] as a classic replicator, dependent on *R*_*k*_ consumption by means of *ξ*_*ik*_ that weighs the strength of the resource-consumer interaction. The terms in *ξ* come from a uniform rectangular distribution U [0, 1], with connectivity *C* = 0.3. The HGT term depends on the transmissibility *τ* of the recombinant vector and the peer-to-peer encounter with a conjugated synthetic donor. Plasmid loss is allowed, therefore *S*_*i*_ is transformed to *N*_*i*_ at rate *µ*. Finally, there is a decay rate, represented by *σ*_*i*_ ∈ [0, 1], and to prevent unrealistically low biomass, a small immigration factor of *ϵ* = 1 *×* 10^−5^ is introduced.

The synthetic strain grows at a rate 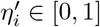, which is assumed to be lower than that of their wild-type counterparts, i.e., 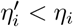, due to the metabolic burden imposed by the recombinant load. The resource-consumer interaction matrix *ξ*_*ik*_ is shared between both wild-type and synthetic species. The HGT term remains the same but contributes positively, since it represents the recruitment of individuals carrying plasmids. In contrast, *µ* denotes the plasmid loss rate. The decay rate *σ*_*i*_ is assumed to be unchanged. Note that there is no immigration term for the synthetic population. While the wild-type community benefits from an established ecological assembly process (with a rate immigration term *ϵ*) the synthetic population *S*_*i*_ originates solely through HGT, at least during the initial phase of ecological engineering.

Finally, the effect encoded in the plasmid is captured by the following function:

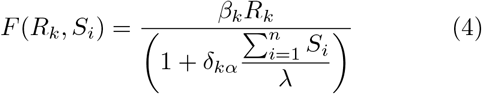

The function *F* (*R*_*k*_, *S*_*i*_) introduces a non-linear adjustment to the resource decay dynamics. Specifically, the existence of synthetic species in the community induces a damping effect on the degradation of a specific resource or a group of resources *α*. The maximum decay rate for a given resource *k* is denoted by *β* ∈ [0, 1], while the constant *λ* ∈ [0, 1] represents the rate at which the degradation process (*β*) is inhibited due to the presence of the synthetic strain. Here *δ*_*kα*_ = 1 when *k* = *α* and zero otherwise. Hence, the damping effect exclusively influences the targeted resource. Any subset of the resources 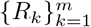 can be targeted (from one to all).

## III. RESULTS

### Low-dimensional model: Predictive Insights

As an initial approach, we analyzed a simplified coarse-grained resource-consumer system Fig. 3a. In this way, we can obtain some relevant insight concerning the expected behaviour of our multidimensional system. One way of coarse-graining the original model is to assume a homogeneous set of parameters, namely that all parameters can be well represented by some average, constant value. In our case that means taking *σ*_*i*_ = *σ, ϵ* = 0 *ξ*_*ij*_ = *ξ*. Additionally, without loss of generality, we take *ξ* = 1. The total values of the variables can be represented as *R* = ∑_*i*_ *R*_*i*_, *N* = ∑_*i*_ *N*_*i*_ and *S* = ∑_*i*_ *S*_*i*_, therefore if we have an aggregated dynamics decribed by:

**FIG. 3.**
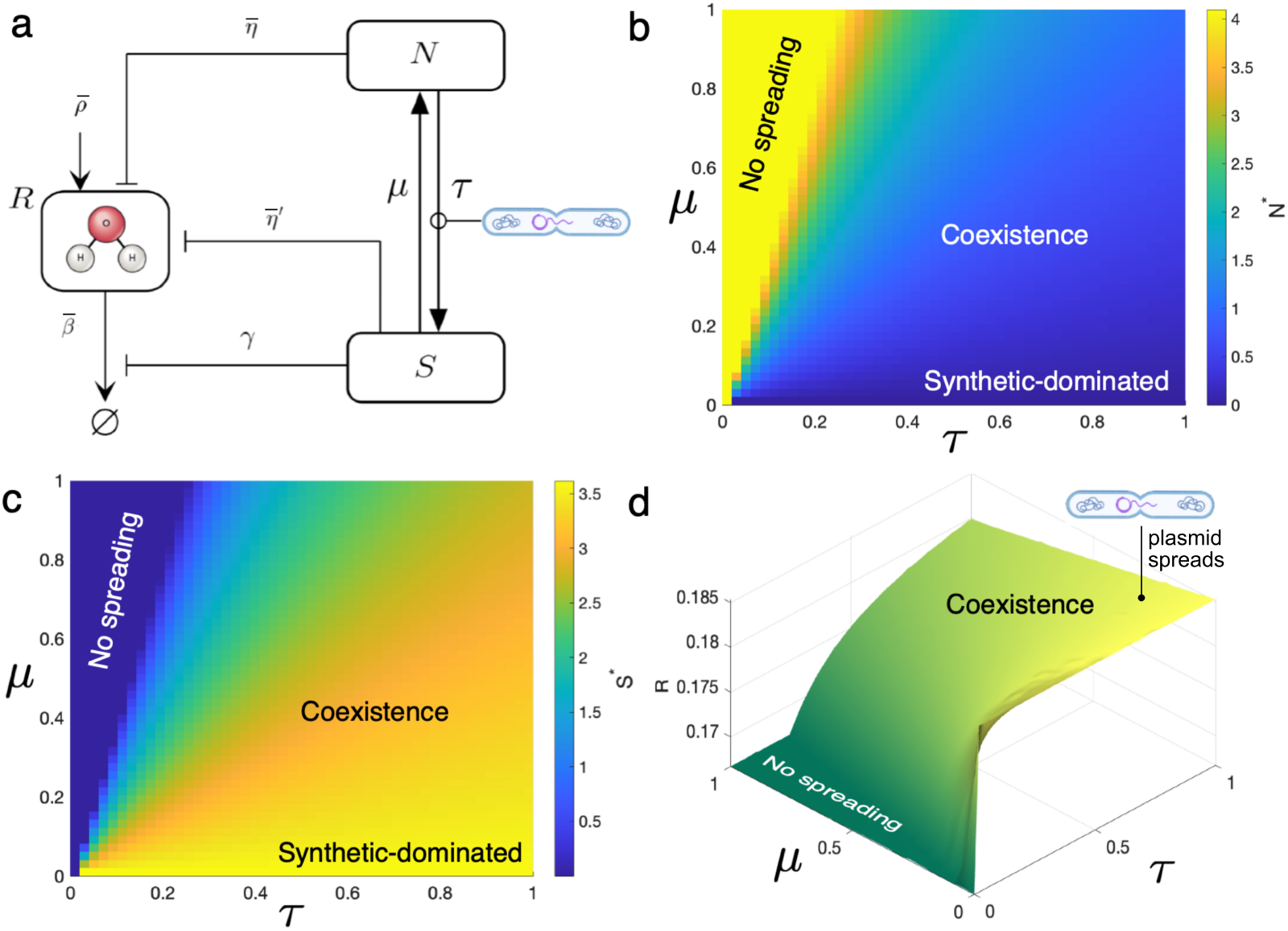
A coarse-grained, low-dimensional model of HGT dynamics. Here, the diverse community structure is reduced to pairwise dynamics (Eqs. 5-7) between a wild-type species *N* and its synthetic counterpart *S*, both competing for a single resource *R*. The results from this model (see SM for details) are shown in (b-d), where we plot the stable populations of (a) wild type, (b) synthetic, and (c) resource levels. Two well-defined phases exist, associated with the failure of transmission and its success. Here (a): *ρ* = 0.8, *β* = 0.7, *γ* = 0.3, *η* = 0.6, *η*^*′*^ = *η* − 0.1*η* and *σ* = 0.1.

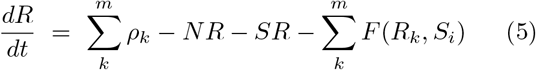

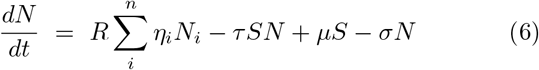

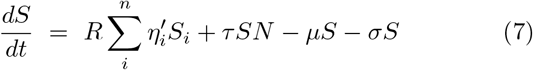

The function 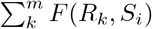 stands for:

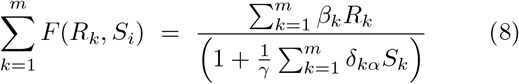

were the *δ*_*kα*_∑ _*i*_ *S*_*i*_ that affect certain resources can be simplified, assuming that all resources are affected. Then *δ*_*kα*_ = 1.

For the rest of the parameters we can use the expected averages as:

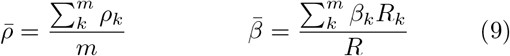

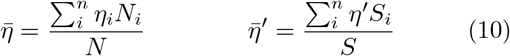

Therefore, it can be shown that the previous equations (1, 2, and 3) collapse in a three-dimensional model, namely:

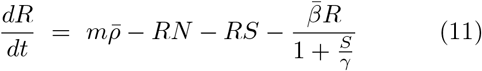

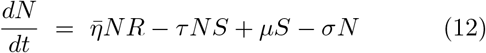

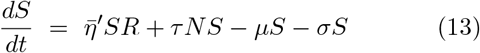

The analysis of this system reveals three fixed points, obtained from *dN/dt* = *dS/dt* = *dR/dt* = 0, namely: (1) resource-only, 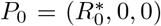, (2) resource and wild type, 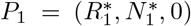 and (3) the coexistence point, 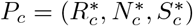. Here we define:

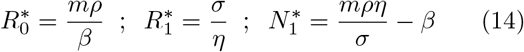

An interesting first result is that none of the attractors contains *N*^*^ = 0 and *S*^*^ *>* 0. This means that, in the mean field approximation, the mutant cannot completely overtake its wild-type counterpart, except for the limit case scenario where *µ* = 0. This shows that natural loss of the engineered plasmid by the synthetic strain is not detrimental but is, in fact, useful to keep the wild-type strain in place.

Furthermore, the stability analysis of the coexistence phase shows that there is a critical boundary separating a domain of successful HGT (when *P*_*c*_ is stable) from the extinction of the synthetic (when *P*_1_ is stable) (see SM). This condition is a linear relation:

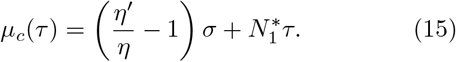

For *µ < µ*_*c*_(*τ*), *S* propagates successfully and for *µ > µ*_*c*_(*τ*), *S* cannot persist.

The basic properties and steady states of this system are summarized in the bifurcation diagrams shown in Fig. 3. Here we can see the equilibrium points for *N*^*^, *S*^*^, and *R*^*^, against different values of two key parameters relevant for our engineering approach, namely the efficiency of the HGT process (*τ*) and the reversal of the process due to loss of plasmids, as weighted by *µ*. As discussed above, the system has two nontrivial stable attractors, involving either HGT failure (only the wild-type survives) or success, whose parameter domains are indicated in Figs. 3b-d as “no spreading” and “coexistence”. As predicted by the theory, a straight line *µ*_*c*_(*τ*) separates them. Interestingly, the region where HGT succeeds does not depend on the engineered function *F* : To ensure that a plasmid and synthetic strain *S* are maintained in a population, we only need to focus on tuning *µ* and *τ* accordingly.

In that regard, and because the implemented function *F* reduces the degradation of the resource, we can see that higher *τ* implies higher *S* and, in turn, more resource (Fig. 3d). However, because this effect saturates at a certain *S*, the model indicates that we do not need to aim towards maximum *S*. Instead, a small presence of *S* can already increase the amount of resource while maintaining the wild-type strain in place (Fig. 3d). This again indicates that moderate values of the spread (*τ*) and loss (*µ*) of the plasmid might be sufficient to achieve the desired effect.

The main message from the low-dimensional model is that, provided that the rate of HGT is high enough to compensate for the plasmid loss rate, we should expect to observe a successful propagation of our engineered plasmid. In this co-existence phase, the engineered functionality would be effective. We next ask if these predictions for co-existence and successful functionality at intermediate *τ* and *µ* values hold against a more realistic scenario, where multiple species and resources are explicitly introduced.

### Multispecies ecosystem

Using Equations (1)–(3), we numerically integrated a high-dimensional consumer–resource system with a fourth-order Runge–Kutta scheme. Each community had *m* = 20 species and *n* = 20 resources and was evolved to stability over Δ*t* = *T*_1_ = 200. At *T*_1_, we introduced an invading synthetic strain *S*_*i*_ derived from a randomly chosen wild type *N*_*i*_ with *N*_*i*_(*T*_1_) *>* 0.2, using an inoculum *S*_*i*_(*T*_1_) = 0.01 (Maull et al., 2024). The system was then simulated for 200 additional steps to a new equilibrium at *T*_2_, allowing up to 20 synthetic counterparts (one per wild type) via HGT. Synthetic strains shared the wildtype interaction matrix; plasmid acquisition imposed a 10% metabolic burden, *η*^*′*^ = 0.9 *η*.

We swept plasmid transmissibility and loss rate, *τ* ∈ [0, 0.15] and *µ* ∈ [0, 0.3]. For each (*τ, µ*) pair, 200 independent replicas were run with randomized parameters *ρ*_*k*_ ∈ [0.8, 1], *β*_*k*_ ∈ [0.7, 0.9], *γ* = 0.3, *η*_*i*_ ∈ [0.5, 1], *σ*_*i*_ ∈ [0.5, 0.7] and *ϵ* = 10^−5^. The resource affected by *S* are determined by *δ*_*i*_, see methods.

Community change from *T*_1_ to *T*_2_ was quantified by: (i) synthetic relative abundance,

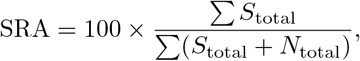

computed over species with abundance *> ϵ* and averaged over the 200 replicas per (*τ, µ*); (ii) strain biodiversity fold change,

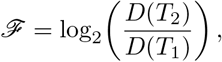

with *D*(*t*) the number of strains (wild-type + synthetic) above *ϵ*; (iii) biomass fold change,

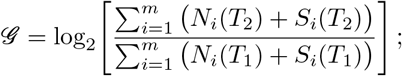

We report fold change on a log_2_ scale. Values *>* 0 indicate an increase, *<* 0 a decrease, and 0 no change. Key benchmarks: ℱ | 𝒢 = 1 (doubling, 2×), ℱ | 𝒢 = − 1 (halving, 1*/*2×), ℱ | 𝒢 = −2 (1*/*4×), etc.

All metrics were averaged over the 200 replicas per (*τ, µ*); detailed results appear in the Supplementary Information.

### Coexistence as the main attractor

A central objective of our multispecies model is to explore how the introduction of an engineered plasmid influences the structure and dynamics of a complex microbial community. This question is closely tied to a long-standing concern in ecological engineering: the risk of negative cascading effects when introducing modified organisms into the wild. As highlighted in (Maull et al., 2024; Maull and Solé, 2022), one proposed strategy to mitigate such risks is to minimize ecological disruption by engineering strains that are derived from wild-type species already present in the community. In the absence of horizontal gene transfer (HGT), this approach appears relatively safe, since the modified strain remains confined to a specific ecological and genetic niche. However, the introduction of HGT as a tool for ecosystem engineering could change the landscape. With HGT, genetically modified material can spread across multiple species boundaries, potentially reshaping ecological networks in ways that are difficult to anticipate. As shown below, the mathematical model strongly points otherwise to a robust community response with no biodiversity loss and improved functionality.

One way to quantify structural changes in the community after HGT is by measuring the Kullback–Leibler (KL) divergence, which captures how much one distribution differs from another, in this case, the relative abundances of strains before and after inoculation. Intuitively, a low KL value means that the overall community composition remains similar. Across simulations, KL values remain consistently low (see SM), suggesting that, although individual biomasses and strain identities may change (see the sections below), the broader structure of the community and the coexistence attractor remain stable. For instance, if the original community was dominated by strains 2, 1, and 3, the new configuration typically retains a similar profile, now possibly dominated by the plasmid-carrying variants 2’, 1’, and 3’ (Fig. 4a).

**FIG. 4.**
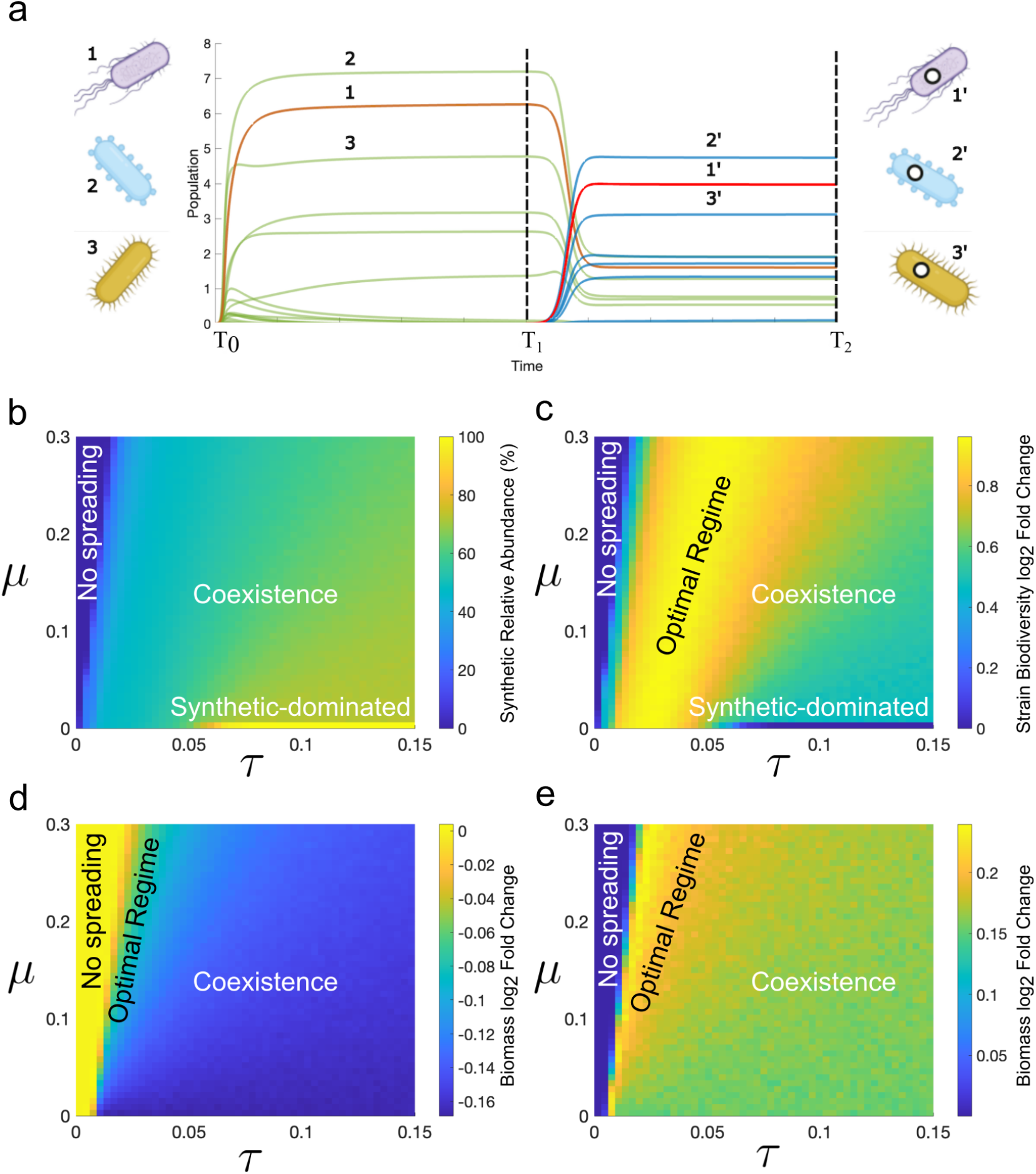
Statistical analysis of the high-dimensional model for *η*^*′*^ = *η* − 0.1*η* for a community with *n* = *m* = 20. After the first inoculation of the synthetic plasmid carrier at *t* = *T*_1_, a new stability state is reached at *t* = *T*_2_. Panel (a) illustrates the assembly process to *T*_1_, with color coding for wild-type (green), synthetic (blue), initial wild-type chosen plasmid carrier (orange), and its synthetic counterpart (red). In (b), the relative abundance of *S*_*i*_ vs *N*_*i*_ and in (c) the biodiversity fold change across *τ* values are shown, both results at *t* = *T*_2_. Panel (d) mirrors (c) but with biomass fold change. Panel (e) also shows biomass fold change at *t* = *T*_2_, but in this case, the totality of the resources is positively affected by the synthetic species, likewise following equation 4. Here above-zero fold change values are observed despite metabolic burden drawbacks. Interestingly, in all cases (b,c, and d), a sweet spot in terms of biodiversity and biomass production appears near the transition point. The results remain consistent across parameters, here with randomized values: *ρ*_*k*_ ∈ [0.8, 1], *β*_*k*_ ∈ [0.7, 0.9], *γ* = 0.3, *η*_*i*_ ∈ [0.5, 1], *η*^*′*^ = *η* − 0.1*η* and *δ*_*i*_ ∈ [0.5, 0.7].

Once again, the coexistence attractor, where both wild-type and synthetic strains are present, dominates the space of possible outcomes (Fig. 4b). Wild-type strains only fully disappear in the extreme case where *µ* = 0 and the plasmid is never lost. Interestingly, in the multispecies context, plasmid spread increases the space of coexistence compared to the low-dimensional model. Because HGT acts as an epidemic process between species, having more potential host species increases the effective transmission rate (see SM). This, in turn, makes the spread of the plasmid easier: Even a small transmission rate *τ* both within and between species quickly ensures the success of the HGT intervention and allows the long-term coexistence of synthetic and wild-type strains.

### Strain Biodiversity Increase

At *τ* = 0, the plasmid cannot propagate, and only wild-type strains persist. In contrast, when *µ* = 0, the plasmid is never lost and synthetic variants fully occupy the system. Between these extremes, we observe varying degrees of coexistence. Here, we ask whether this coexistence increases the overall biodiversity of the strain, that is, the total number of distinct strains, or whether some species are lost in the process.

A central result of our analysis is that, if *m*^*^ denotes the number of wild-type species that survive in steady state before inoculation of the plasmid, then the total biodiversity of the system is always equal to or greater than *m*^*^. In fact, biodiversity can reach 2*m*^*^, resulting in a two-fold increase in strain diversity (Fig. 4c; see Methods and SM). At the two limiting cases (*τ* = 0 or *µ* = 0), the community contains either *m*^*^ wild-type strains or *m*^*^ synthetic variants, respectively. However, in the intermediate regime, we consistently find more than *m*^*^ strains, indicating that some species persist in both wildtype and synthetic forms.

At high transmissibility (*τ*), all synthetic strains are present, along with a subset of the most abundant wild types. In this case, rarer wild-type strains are typically replaced by their plasmid-carrying counterparts (see SM for a detailed analysis). In contrast, at intermediate values of *τ*, the system reaches maximum strain diversity, with all species maintaining both wild-type and synthetic forms. As predicted by the low-dimensional model, this result shows that a high *τ* is not required for a success-ful HGT intervention. Instead, intermediate values of *τ* define an optimal regime, where synthetic strains are effectively introduced without displacing their wild-type counterparts.

### Total Biomass Gain

Synthetic variants carry an inherent trade-off: While they can reduce resource degradation (parameterized by *γ*), they are also less efficient consumers, with *η*^*′*^ *< η*. This raises a key ecological question: Does the introduction of synthetic strains lead to a net increase in total community biomass due to improved resource conservation, or does it reduce biomass as a result of diminished replication efficiency? To investigate this, we quantify the fold change in total biomass from time *T*_1_ to time *T*_2_, as described in the Materials and Methods section. This metric captures the net impact of engineering interventions on community productivity.

Our analysis reveals a clear pattern. When the engineered plasmid targets a single resource, the biomass fold change is generally negative across most of the coexistence region (Fig. 4d), indicating a net loss in total biomass. This suggests that, under this minimal engineering scenario, the cost of carrying the plasmid outweighs its ecological benefit. This outcome aligns with a conservative baseline assumption, where the metabolic burden imposed by reduced efficiency dominates over the limited savings in resources. However, there is a notable exception: Near the optimal parameter regime discussed earlier, the biomass loss is less pronounced. This is visible in Fig. 4d as light green areas, where total biomass remains close to or slightly above baseline levels. In these cases, most individuals in the community remain wildtype, and only a small fraction carries the synthetic plasmid, minimizing the collective cost.

But what if the engineered function affects multiple resources simultaneously? Consider, for example, encoding a gene that synthesizes a hygroscopic molecule such as hyaluronic acid, which enhances water retention in the environment. This could not only improve access to water, but also indirectly improve the availability of several other resources (Li et al., 2001; Sze et al., 2016; Tamaru et al., 2005; Widner et al., 2005). This multiresource enhancement scenario is explored in detail in the Supplementary Materials, where we systematically vary the number of resources affected by the engineered function, from one to all.

As shown in Fig. 4e, when all resources are positively impacted by the engineered strains, the change in biomass fold is consistently positive across the entire coexistence region. This indicates a robust ecological benefit of the intervention: even at low values of plasmid transmission rate *τ*, the engineered function promotes higher total biomass without compromising species coexistence. This result underscores the potential of synthetic strategies that enhance broad ecological functions rather than narrowly targeting individual resources.

Importantly, the previously identified optimal region reemerges clearly under this expanded scenario, reaffirming its role as a balance point between metabolic cost and community-wide benefit. In addition, the strain diversity remains stable regardless of the number of resources targeted, as it is primarily governed by the dynamics of horizontal plasmid transfer, a result consistent with both the low-dimensional model and the full simulations discussed above.

### Optimal regime

The presence of an optimal regime is not an obvious feature and warrants careful consideration. The key to understanding it is twofold. From the perspective of the pure dynamics of plasmid spreading, there is a peak in strain biodiversity when the ratio *τ/µ* is low. This scenario maintains a proportionally small number of synthetic individuals in the community; therefore, the metabolic burden, given by *η*^*′*^ = *η* − 0.1*η*, has a reduced impact on the general community, and low-abundance species, which are more susceptible, are less stressed.

Secondly, when the number of affected resources is sufficiently high, biomass gain is optimized by a synthetic community that is small enough to avoid triggering a global loss due to metabolic burden, yet large enough to produce a meaningful improvement in resource dampening. This trade-off is clearly evident in Equation 4. Resources are depleted through both consumption and natural decay, as described in Equation 1. Importantly, the decay term in Equation 4 is modulated by the presence of the synthetic community, introducing a dampening dynamic. If we evaluate the following limit:

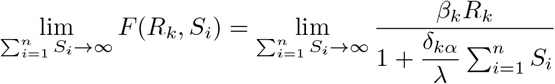

Observe that as 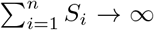, or the synthetic community reaches its maximum size, the denominator behaves as:

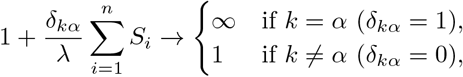

Therefore, the limit becomes:

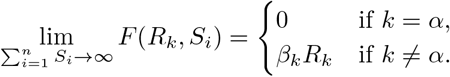

When a resource is affected, its depletion cannot fall below its consumption rate, leading to a plateau in potential gain. Expanding the synthetic community beyond a certain point does not proportionally increase the positive effect on resource availability. Instead, an optimal balance, or optimal regime, can be achieved by tuning, through design, the plasmid’s transmissibility and its loss potential (Aparicio et al., 2022).

## IV. DISCUSSION

Could horizontal gene transfer be harnessed as a safe and effective tool for ecosystem restoration? Our results suggest that the answer is yes: When applied with precision, synthetic biology could enhance and stabilize microbial communities through horizontal gene transfer (HGT) without affecting ecological balance. Inspired by soil microbiomes as a general case study, we demonstrate that engineered strains carrying functional plasmids can be successfully spread through microbial networks under controlled conditions. This strategy not only preserves strain-level biodiversity but can, in some cases, also lead to a net increase in total community biomass.

A simplified analytical model revealed that the balance between plasmid transmissibility (*τ*) and loss rate (*µ*) determines whether synthetic strains can successfully establish within a community. When *τ/µ* exceeds a critical threshold, coexistence between synthetic and wildtype strains becomes the dominant outcome, a result that aligns well with the behavior observed in more detailed, multidimensional simulations.

Importantly, using native species as initial plasmid carriers ensures that synthetic strains remain ecologically compatible, occupying the same niche as their wildtype counterparts. This design principle prevents major changes in community composition, as confirmed by negligible Kullback–Leibler divergence before and after synthetic introduction. Coexistence emerges as the primary attractor in a wide range of conditions, with plasmid spread limited by loss rates and niche overlap, thereby avoiding disruptive dominance by engineered variants. In practice, this suggests not maximal spread, but rather constructs whose conjugation range and frequency are sufficient to offset realistic plasmid loss.

HGT systematically increases strain-level biodiversity (Fig. 4c). Each successful transfer adds a synthetic counterpart, usually without displacing its original wild-type ancestors. This diversification peaks within an “optimal region” in parameter space, where propagation is efficient but controlled, and the metabolic cost of plasmid carriage remains modest. As a result, the modest and persistent presence of the plasmid is often preferable to aggressive spread. And it is sufficient to elevate resources and diversity while avoiding unnecessary costs.

Function targeting is pivotal for success. When the engineered cassette buffers only a single limiting resource, biomass responses are modest or even slightly negative, as the metabolic burden of carrying the new plasmid offsets the ecological benefit. In contrast, when the cassette acts as a *keystone gene*, that is, one whose expres-sion simultaneously affects multiple limiting factors, the effects become pronounced at the community level. Hygroscopic exopolymers provide a concrete example: By retaining water, they indirectly stabilize several resources at once, shifting the system toward higher biomass across a broad coexistence region (Fig. 4e). More generally, engineered functions that couple to systemic constraints, such as hydrology, nutrient retention, or pH buffering, are likely to outperform narrowly targeted constructs, as they relieve integrated ecological bottlenecks rather than isolated ones.

These properties also have safety implications. First, because the coexistence phase is a generic outcome of the model, wild-type strains tend to persist unless plasmid loss becomes negligible (*µ* → 0). As a result, complete replacement of wild types is unlikely under realistic conditions. Second, the saturating form of the function *F* imposes diminishing returns: increasing transmissibility beyond the optimal range provides little additional benefit while increasing the metabolic burden. Furthermore, the functional form of *F* captures the decay-damping effect on a targeted resource, which represents one of the most stringent scenarios. Other functional mechanisms, such as detoxification, post-fire alkalinity buffering, or heavy metal remediation, may couple differently to resource dynamics and should be examined in future studies.

In summary, by coupling consumer-resource dynamics with the epidemic-like spreading of a synthetic gene, our framework reveals the feasibility of ecological engineering through synthetic HGT. Our results inform design strategies by proposing guiding principles for the selection of appropriate gene flow rates and ecological functions. Future work should explore spatial heterogeneity, evolutionary feedbacks, and experimental validation to assess long-term viability. These findings offer a concrete design roadmap for the deployment of engineered microbes in support of ecosystem resilience and restoration.

## Acknowledgments

VM and RS thank Jordi Piñero, Luis Seoane, and Jordi Pla for fruitful discussions. GA-G thanks Sonia Kéfi for her support. Special thanks to Maryam Abu Daqa and Ruwaida Kamal Amer for inspiration. VM is supported by the Ajuntament de Barcelona and “la Caixa” Foundation. RS is supported by the AGAUR 2021 SGR 0075 grant. GA-G is supported by a Marie Sklodowska-Curie Actions Postdoctoral Fellowship under project FRAGILEPRINTS 101105029. Views and opinions expressed are, however, those of the author(s) only and do not necessarily reflect those of the European Union or the CNRS. Neither the European Union nor the CNRS can be held responsible for them. The authors also appreciate the support of the Santa Fe Institute, where most of the work was done.

## Notes

### Competing Interest Statement

The authors have declared no competing interest.

